# Exploring systematic biases, rooting methods and morphological evidence to unravel the evolutionary history of the genus *Ficus* (Moraceae)

**DOI:** 10.1101/2020.04.15.042259

**Authors:** Jean-Yves Rasplus, Lillian Jennifer Rodriguez, Laure Sauné, Yang-Qiong Peng, Anthony Bain, Finn Kjellberg, Rhett D. Harrison, Rodrigo A.S. Pereira, Rosichon Ubaidillah, Christine Tollon-Cordet, Mathieu Gautier, Jean-Pierre Rossi, Astrid Cruaud

**Affiliations:** CBGP, INRAE, CIRAD, IRD, Montpellier SupAgro, Univ Montpellier, Montpellier, France; Institute of Biology, University of the Philippines Diliman, Quezon City, Philippines; Natural Sciences Research Institute, University of the Philippines Diliman, Quezon City, Philippines; CAS Key Laboratory of Tropical Forest Ecology, Xishuangbanna Tropical Botanical Garden, Chinese Academy of Sciences, Kunming, China; Department of Biological Sciences, National Sun Yat-sen University, Kaohsiung, Taiwan; CEFE, CNRS—Université de Montpellier—Université Paul-Valéry Montpellier—EPHE, Montpellier, France; World Agroforestry, Eastern and Southern Africa, Region, 13 Elm Road, Woodlands, Lusaka, 10101 Zambia; Departamento de Biologia, FFCLRP, Universidade de São Paulo, Ribeirão Preto, SP, Brazil; Museum Zoologicum Bogoriense, LIPI, Gedung Widyasatwaloka, Jln Raya km 46, Cibinong, Bogor 16911, Indonesia; AGAP, INRAE, CIRAD, Montpellier SupAgro, Univ Montpellier, Montpellier, France

**Keywords:** compositional bias, fig trees, long branch attraction, morphology, phylogeny, RAD-seq, traits

## Abstract

Despite their ecological and evolutionary importance as key components of tropical ecosystems, the phylogeny of fig trees is still unresolved. We use restriction-site-associated DNA (RAD) sequencing (*ca* 420kb) and 102 morphological characters to elucidate the relationships between 70 species of *Ficus* representing all known subgenera and sections and five outgroups. We compare morphological and molecular results to highlight discrepancies and reveal possible inference bias. We analyse marker and taxon properties that may bias molecular inferences, with existing softwares and a new approach based on iterative principal component analysis to reduce variance between clusters of samples. For the first time, with both molecular and morphological data, we recover a monophyletic subgenus *Urostigma* and a clade with all gynodioecious fig trees. However, our analyses show that it is not possible to homogenize evolutionary rates and GC content for all taxa prior to phylogenetic inference and that four competing positions for the root of the molecular tree are possible. The placement of the long-branched section *Pharmacosycea* as sister to all other fig trees is not supported by morphological data and considered as a result of a long branch attraction artefact to the outgroups. Regarding morphological features and indirect evidence from the pollinator tree of life, the topology that divides the genus *Ficus* into monoecious *versus* gynodioecious species appears most likely. Active pollination is inferred as the ancestral state for all topologies, ambiguity remains for ancestral breeding system including for the favored topology, and it appears most likely that the ancestor of fig trees was a freestanding tree. Increasing sampling may improve results and would be at least as relevant as maximizing the number of sequenced regions given the strong heterogeneity in evolutionary rates, and to a lesser extent, base composition among species. Despite morphological plasticity and frequent homoplasy of multiple characters, we advocate giving a central role to morphology in our understanding of the evolution of *Ficus*, especially as it can help detect insidious systematic errors that tend to become more pronounced with larger molecular data sets.

## INTRODUCTION

*Ficus* (Moraceae) is a pantropical and hyperdiverse genus (*ca* 850 species) that includes a broad range of growth forms (trees, hemi-epiphytes, shrubs, climbers) with diverse ecologies (Berg and Corner 2005, Harrison 2005, Harrison and Shanahan 2005). As their inflorescences (figs) are important food source for hundreds of frugivorous species (Shanahan et al. 2001), fig trees are key components of tropical ecosystems. They are also known for their intricate relationships with their pollinating wasps (Agaonidae). Indeed, since *ca* 75 Myr, fig trees and agaonids have been obligate mutualists (Cruaud et al. 2012). The wasp provides pollination services to the fig tree, while the fig tree provides breeding sites for the wasps, and none of the partners are able to reproduce without the other (Galil 1977, Cook and Rasplus 2003). Fifty-two percent of *Ficus* species are monoecious, while 48% are gynodioecious. In monoecious species, figs contain staminate and pistillate flowers and produce pollen, pollinators and seeds. Gynodioecious species are functionally dioecious with male function (pollen and pollinator production), and female function (seed production) segregated on separate individuals. *Ficus* species are pollinated either actively (two-thirds) or passively (one-third) (Kjellberg et al. 2001). In passively pollinated figs, emerging wasps are dusted with pollen before leaving their natal fig. In actively pollinated figs, wasps use their legs to collect pollen they will later deposit on flowers of receptive figs, while laying their eggs.

Despite its ecological importance, the evolutionary history of the genus remains unclear. Several studies have attempted to reconstruct the phylogeny of *Ficus* using Sanger sequencing of chloroplast markers (Herre et al. 1996), external and/or internal transcribed spacers (ETS, ITS) (Weiblen 2000, Jousselin et al. 2003), a combination of nuclear markers (Rønsted et al. 2005, Rønsted et al. 2008, Xu et al. 2011, Cruaud et al. 2012, Pederneiras et al. 2018, Zhang et al. 2018, Clement et al. 2020) or next-generation sequencing of chloroplast genomes (Bruun-Lund et al. 2017). None of these studies successfully resolved the backbone of the phylogeny and there is no consensus on the relationships between major groups of *Ficus* yet. As a consequence, current classification remains inconsistent with different phylogenetic levels classified under the same taxonomic rank or, to the opposite, identical taxonomic rank appearing at different phylogenetic level (Table 1). In addition, our ability to analyse the evolution of key traits that may have contributed to the evolutionary and ecological success of the genus is limited. Areas of agreement are (i) the position of sect. *Pharmacosycea* as sister to all other fig trees (with poor or high support depending on the study), (ii) a monophyletic and strongly supported *Mixtiflores* (*i.e.* subgenus *Urostigma* excluding sect. *Urostigma, sensu* Clement *et al.* (2020)), and (iii) a monophyletic clade formed by subg. *Synoecia*, sect. *Eriosycea* and subsect. *Frutescentiae*, sometimes recovered sister to subg. *Sycidium*. However, these two last results were challenged by plastome analysis (Bruun-Lund et al. 2017) that provided improved resolution but highlighted a high level of cyto-nuclear discordance with some subgenera undoubtedly monophyletic (e.g. *Sycidium*) recovered as polyphyletic. Non-ambiguous cases of cyto-nuclear discordance were previously detected in African fig trees (Renoult et al. 2009) and are more frequently reported with the use of high throughput sequencing approaches (Huang et al. 2014). Hence, the use of nuclear, genome-wide markers appears more appropriate to resolve the phylogeny of fig trees.

**Table 1.**
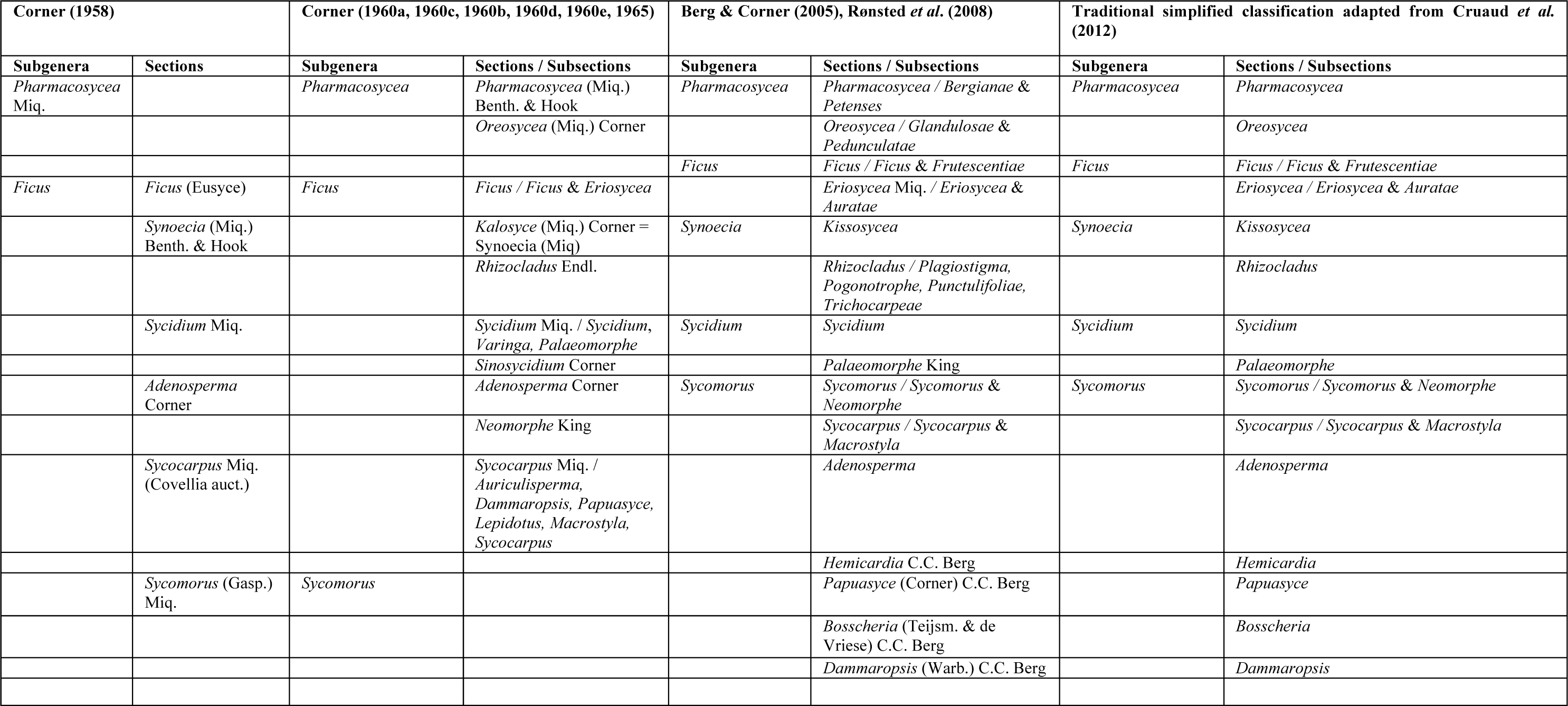
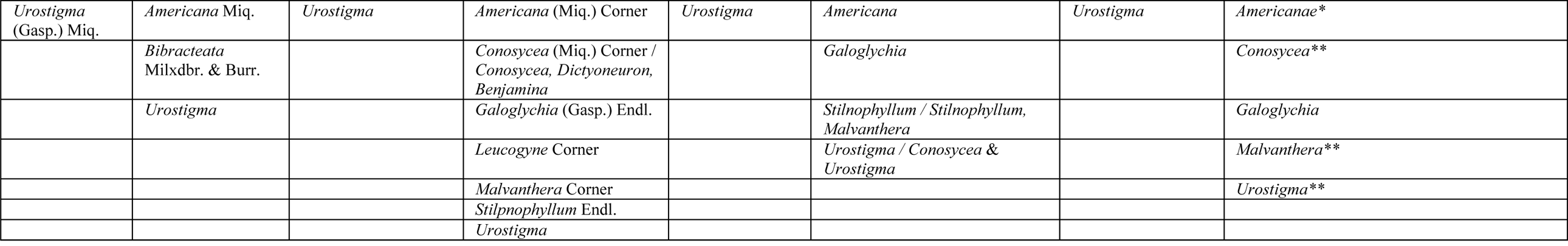
Classification of the genus *Ficus.* In this paper, we use the simplified classification adapted from Cruaud *et al.* 2012. *the proper spelling of this section should be *Americanae*, which is the original spelling proposed by Miquel. Indeed, it is a plural adjective. ** *Conosycea, Malvanthera* and *Urostigma* are considered as sections instead of subsections and *F. elastica* (the only member of the subsect. *Stilnophyllum sensu* Berg & Corner (2005) is included in *Conosycea* as recovered in previous molecular studies. *Nota:* Subgenus *Urostigma* excluding section *Urostigma* is referred to as *Mixtiflores* by Clement *et al.* (2020).

| Corner (1958) |  | Corner (1960a, 1960c, 1960b, 1960d, 1960e, 1965) |  | Berg & Corner (2005), Ronsted *et al.* (2008) |  | Traditional simplified classification adapted from Cruaud *et al.* (2012) |  |
| --- | --- | --- | --- | --- | --- | --- | --- |
| Subgenera | Sections | Subgenera | Sections / Subsections | Subgenera | Sections / Subsections | Subgenera | Sections / Subsections |
| *Pharmacosycea* Miq. |  | *Pharmacosycea* | *Pharmacosycea* (Miq.) Benth. & Hook | *Pharmacosycea* | *Pharmacosycea* / *Bergianae* & *Petenses* | *Pharmacosycea* | *Pharmacosycea* |
|  |  |  | *Oreosycea* (Miq.) Corner |  | *Oreosycea* / *Glandulosae* & *Pedunculatae* |  | *Oreosycea* |
|  |  |  |  | *Ficus* | *Ficus* / *Ficus* & *Frutescentiae* | *Ficus* | *Ficus* / *Ficus* & *Frutescentiae* |
| *Ficus* | *Ficus* (Eusyce) | *Ficus* | *Ficus* / *Ficus* & *Eriosycea* |  | *Eriosycea* Miq. / *Eriosycea* & *Auratae* |  | *Eriosycea* / *Eriosycea* & *Auratae* |
|  | *Synoecia* (Miq.) Benth. & Hook |  | *Kalosyce* (Miq.) Corner = *Synoecia* (Miq.) | *Synoecia* | *Kissosycea* | *Synoecia* | *Kissosycea* |
|  |  |  | *Rhizocladus* Endl. |  | *Rhizocladus* / *Plagiostigma*, *Pogonotrophe*, *Punctulifoliae*, *Trichocarpeae* |  | *Rhizocladus* |
|  | *Sycidium* Miq. |  | *Sycidium* Miq. / *Sycidium*, *Varinga*, *Palaeomorphe* | *Sycidium* | *Sycidium* | *Sycidium* | *Sycidium* |
|  |  |  | *Sinosycidium* Corner |  | *Palaeomorphe* King |  | *Palaeomorphe* |
|  | *Adenosperma* Corner |  | *Adenosperma* Corner | *Sycomorus* | *Sycomorus* / *Sycomorus* & *Neomorphe* | *Sycomorus* | *Sycomorus* / *Sycomorus* & *Neomorphe* |
|  |  |  | *Neomorphe* King |  | *Sycocarpus* / *Sycocarpus* & *Macrostyla* |  | *Sycocarpus* / *Sycocarpus* & *Macrostyla* |
|  | *Sycocarpus* Miq. (Covellia auct.) |  | *Sycocarpus* Miq. / *Auriculisperma*, *Dammaropsis*, *Papuasyce*, *Lepidotus*, *Macrostyla*, *Sycocarpus* |  | *Adenosperma* |  | *Adenosperma* |
|  |  |  |  |  | *Hemicardia* C.C. Berg |  | *Hemicardia* |
|  | *Sycomorus* (Gasp.) Miq. | *Sycomorus* |  |  | *Papuasyce* (Corner) C.C. Berg |  | *Papuasyce* |
|  |  |  |  |  | *Bosscheria* (Teijsm. & de Vriese) C.C. Berg |  | *Bosscheria* |
|  |  |  |  |  | *Dammaropsis* (Warb.) C.C. Berg |  | *Dammaropsis* |
| *Urostigma*<br>(Gasp.) Miq. | *Americana* Miq. | *Urostigma* | *Americana* (Miq.) Corner | *Urostigma* | *Americana* | *Urostigma* | *Americanae* * |
|  | *Bibracteata*<br>Milxdbr. & Burr. |  | *Conosycea* (Miq.) Corner /<br>*Conosycea*, *Dictyoneuron*,<br>*Benjamina* |  | *Galoglychia* |  | *Conosycea* ** |
|  | *Urostigma* |  | *Galoglychia* (Gasp.) Endl. |  | *Stilnophyllum* / *Stilnophyllum*,<br>*Malvanthera* |  | *Galoglychia* |
|  |  |  | *Leucogyne* Corner |  | *Urostigma* / *Conosycea* &<br>*Urostigma* |  | *Malvanthera* ** |
|  |  |  | *Malvanthera* Corner |  |  |  | *Urostigma* ** |
|  |  |  | *Stilpnophyllum* Endl. |  |  |  |  |
|  |  |  | *Urostigma* |  |  |  |  |

Weiblen (2000) proposed the only phylogenetic hypothesis for the genus (46 species) based on morphological data, with a special focus on gynodioecious fig trees. Representatives of all subgenera of *Ficus* and all sections except one (*Galoglychia*) were scored for 61 characters. No outgroups were included and trees were rooted in agreement with the only molecular study available at that time that included only three species of *Ficus* and eight outgroups (Herre et al. 1996). The consensus tree was poorly resolved and some groups were recovered as paraphyletic but this study represents a good starting point. As underlined by Clement & Weiblen (2009), botanists have long noticed high levels of homoplasy in Moraceae, which complicates the assembly of morphological matrices. Besides, matrix assembly requires a thorough knowledge of the target group and scoring characters on a representative number of specimens is time consuming. This certainly explains the shift towards molecular approaches, which is the general trend in systematics, especially since the advent of high-throughput sequencing technologies.

However, overconfidence in molecular data is risky. Indeed, while genome-scale data may contribute to better resolve phylogenetic relationships (Philippe et al. 2005), they can also infer incorrect yet highly supported topologies due to the failure of current models /methods to capture the full complexity of evolutionary processes (systematic error; Swofford et al. 1996, Phillips et al. 2004, Kumar et al. 2012). This is why morphological approaches, even though considered old-fashioned or supposed to be difficult to conceptualize and to interpret, are still crucial in a time of big data overflow (Wiens 2004, Giribet 2015, Wipfler et al. 2016). Indeed, they constitute an independent set of characters which can help to detect errors in inferences based on molecular data.

Here we infer the evolutionary history of 70 species of *Ficus* representing all known subgenera and sections and five outgroups from i) 102 morphological characters and ii) Restriction-site-Associated DNA sequencing (RAD-seq). RAD-seq has been used successfully to infer recent and ancient evolutionary histories of groups of plants (e.g. (Eaton and Ree 2013, Hipp et al. 2014, Hipp et al. 2019) including a community of 11 Panamanian strangler figs from the section *Americanae* (Satler et al. 2019). In a preliminary study, we targeted conserved regions with an infrequent 8-cutter restriction enzyme (*SbfI*) and highlighted the power of RAD-seq to infer deep relationships between fig trees (Rasplus et al. 2018). However, only forward reads were analysed, taxonomic sampling was reduced and a single outgroup with a high level of missing data was used. Results were encouraging but not supported enough to draw definitive conclusions. Here, we go one step further, by increasing taxon sampling and the length of analysed loci. To mine RAD loci into distant outgroup genomes, we assembled paired reads into long loci. This enabled us to decrease missing data associated with the loss of restriction sites with time, while increasing phylogenetic signal for the ingroup. We critically compare morphological and molecular results and discuss similarities and discrepancies. In addition, we performed a thorough analysis of marker and taxon properties that could bias molecular inferences (heterogeneity in base composition and variable evolutionary rates) with existing software and a new approach based on iterative principal component analysis (PCA) to reduce variance between clusters of samples. We also compared the impact of different rooting strategies on molecular tree topology. Finally, we use our data set to revisit the evolution of key traits (life form, breeding system and pollination mode) that may have contributed to the evolutionary and ecological success of the genus.

## MATERIALS AND METHODS

### Sampling and classification

Here we use the classification by Berg & Corner (2005) with some modifications used in Cruaud *et al.* 2012 (Table 1). Seventy species of *Ficus* representing all known subgenera and sections as well as four outgroups were included in the analysis (Table S1). The same individual was used for molecular and morphological studies. Plants, twigs, leaves and figs were photographied before sampling of a few leaves that were dried for molecular purposes. Voucher specimens are archived at CBGP, Montpellier.

### Morphological Data

Species were scored for 102 morphological characters (Appendix S1). Seventy-eight were extracted from earlier phylogenetic studies (Weiblen 2000, Clement and Weiblen 2009, Chantarasuwan et al. 2015) and sometimes redefined, while 24 characters were used for the first time. Whenever possible we cross-validated our observations and accounted for polymorphism using descriptions available in the literature (Corner 1938, Corner 1967, Corner 1969b, Corner 1969a, Corner 1970, Corner 1978a, Corner 1978b, Berg and Wiebes 1992, Berg and Corner 2005, Berg 2009, Berg et al. 2011) and conspecific specimens from other localities. Data were analysed using Maximum Parsimony (MP) as implemented in PAUP* version 4.0a (Swofford 2003). We used a heuristic search with 5000 random addition sequences (RAS) to obtain an initial tree and “tree bisection and reconnection (TBR)” as branch swapping option, with reconnection limit set to 100. One tree was retained at each step. Characters were equally weighted and treated as unordered and non-additive. Multiple states were interpreted as polymorphism and gaps (characters that were impossible to score because the feature was non-existent) were treated as missing data. Robustness of the topology was assessed by bootstrap procedures (100 replicates; TBR RAS 100; one tree retained at each step). Character transformations were mapped on the majority-rule consensus tree and the four alternative RAD topologies in PAUP* using the ACCTRAN optimization strategy.

### DNA extraction and library construction

Leaves were either dried with silica gel or sun-dried. Twenty mg of dried leaves were placed in Eppendorf vials and crushed with ceramic beads in liquid nitrogen. DNA was extracted with the Chemagic DNA Plant Kit (Perkin Elmer Chemagen, Baesweller, DE, Part # CMG-194), according to the manufacturer’s instructions with a modification of the cell lysis. The protocol was adapted to the use of the KingFisher Flex™ (Thermo Fisher Scientific, Waltham, MA, USA) automated DNA purification workstation. The powder was suspended in 400uL Lysis buffer (200mM Tris pH = 8.0, 50mM EDTA, 500mM NaCl, 1.25 % SDS, 0,5 % CTAB 1% PVP 40000, 1 g/100ml Sodium Bisulfite) and incubated 20 min at 65°C. Then 150 µL of cold precipitation buffer (sodium acetate 3M, pH 5.2) was added. Samples were centrifuged 10 min at 12000 rpm and 350 µL of the supernatant were transferred in a 96 deepwell plate. Binding of DNA on magnetic beads, wash buffer use and elution of purified DNA followed Chemagic kit protocol and KingFisher Flex use.

Library construction followed Baird *et al.* (2008) and Etter *et al.* (2011) with modifications detailed in Cruaud *et al.* (2014) and below. To infer deep phylogenetic relationships, we targeted conserved regions with an infrequent 8-cutter restriction enzyme (*SbfI*). The expected number of cut sites was estimated with the radcounter_v4.xls spread sheet available from the UK RAD Sequencing Wiki (www.wiki.ed.ac.uk/display/RADSequencing/Home). We assumed a 704 Mb approximate genome size (Ohri and Khoshoo 1987), ca 1.44 pg on average for 15 species of *Ficus*) and a 48% GC content (estimated from EST data available on NCBI). Based on those estimates, 9,095 cut sites were expected. 125ng of input DNA was used for each sample. After digestion, 1 uL of P1 adapters (100nM) was added to saturate restriction sites. Samples were then pooled sixteen by sixteen and DNA of each pool was sheared to a mean size of *ca* 400 bp using the Bioruptor® Pico (Diagenode) (15sec ON / 90sec OFF for 8 cycles). After shearing, end repair and 3’-end adenylation, DNA of each pool was tagged with a different barcoded P2 adapter. A PCR enrichment step was performed prior to KAPA quantification. The 2*125nt paired-end sequencing of the library was performed at MGX-Montpellier GenomiX on one lane of an Illumina HiSeq 2500 flow cell.

### Data cleaning and assembly of paired reads into RAD loci

Data cleaning was performed with RADIS (Cruaud et al. 2016), which relies on Stacks (Catchen et al. 2013) for demultiplexing and removal of PCR duplicates. Individual loci were built using *ustacks* [m=15; M=2, N=4; with removal (r) and deleveraging (d) algorithms enabled]. The parameter n of *cstacks* (number of mismatches allowed between sample loci when building the catalog) was set to 20 to cluster enough loci for the outgroups, while ensuring not to cluster paralogs in the ingroup. To target loci with slow or moderate substitution rate, only loci present in 75% of the samples were analysed. Loci for which samples had three or more sequences were removed from the analysis. Loci were aligned with MAFFT v7.245 (-linsi option) (Katoh and Standley 2013). The 583 loci obtained in this step (mergeR1 dataset) were used as a starting point to assemble paired reads into longer RAD loci. The pipeline, scripts and parameters used for the assembly of paired reads are available from https://github.com/acruaud/radseq_ficus_2020. Briefly, for each sample, forward reads used to build the 583 cstacks loci of the mergeR1 data set as well as corresponding reverse reads were retrieved from original fastq files with custom scripts and assembled with Trinity (Haas et al. 2013). Contigs were aligned to the reference genome of *F. carica* assembly <u>GCA_002002945.1</u> with Lastz Release 1.02.00 (Harris 2007). Homology between cstacks loci and reference genome and homology between sample contigs within cstacks loci were tested as follows. For each cstacks locus, the genome scaffold with the highest number of alignment hits was considered as likely to contain the RAD locus. When contigs aligned with different parts of the same scaffold, the genome region that showed the highest identity with the sample contigs (as estimated with Geneious 11.1.4: https://www.geneious.com) was considered as the most likely RAD locus. Cstacks loci for which the sample of *F. carica* JRAS06927_0001 included in the RAD library was not properly aligned with the reference genome (hard or soft clipped unaligned ends > 10 bp) or for which the majority of contigs did not align with the same genome region as *F. carica* JRAS06927_0001 were removed. Finally, loci for which at least one sample contig appeared more than once were discarded. When several contigs were retained per sample (e.g when forward and reverse reads did not overlap; or in case of polyploidy or sequencing mistake) a consensus was built and the IUPAC code was used without considering any threshold, if, for a given position, different nucleotides were present. Of the 583 initial loci, 530 successfully passed quality controls and were retained for phylogenetic analysis.

### Retrieval of RAD loci in genome of outgroup species

As expected for a RAD experiment (Rubin et al. 2012, Gautier et al. 2013), outgroups included in the library had a high level of missing data (*ca* 70%). To decrease missing data, we mined RAD loci in available outgroup genomes. Aside from *F. carica*, only two genomes of Moraceae were available on NCBI when we performed this study: *Morus notabilis* (assembly <u>GCA_000414095.2</u>) and *Artocarpus camansi* (assembly <u>GCA_002024485.1</u>). Pipeline, scripts and parameters used for the retrieval of RAD loci in genome of outgroups are available from https://github.com/acruaud/radseq_ficus_2020. Briefly, the 530 RAD loci extracted from the genome of *F. carica* in the previous step were aligned with the two outgroups genomes using Lastz. Alignment results were parsed with Samtools (Li et al. 2009) to select among the genome regions on which a single RAD locus matched. We considered that the genome region with the highest similarity was the most likely to be homologous with the query RAD locus. Putative RAD loci were extracted from the genome with custom scripts and aligned with the contigs of forward and reverse reads produced in the previous step using MAFFT v7.245 (-linsi option). The final dataset (mergeR1R2) was composed of 530 loci, 71 ingroup species (70 included in the RAD library plus the genome of *F. carica*) and five outgroup species (*Antiaris toxicaria, Artocarpus sp.* and *Morus alba* that were included in the RAD library plus the genomes of *Artocarpus camansi* and *Morus notabilis*). Summary statistics for data sets and samples were calculated using AMAS (Borowiec 2016). Test for phylogenetic signal of sample properties were conducted in R (R Core Team 2018) using the K statistic (Blomberg et al. 2003) implemented in the package Phytools (Revell 2012). The null expectation of K under no phylogenetic signal was generated by randomly shuffling the tips of the phylogeny 1000 times.

### Data set cleaning

To reduce inference bias due to possible misalignment or default of homology, TreeShrink (Mai and Mirarab 2018) was used to detect and remove abnormally long branches in individual gene trees of the mergeR1R2 dataset. As suggested in the manual, the value of b was determined by a preliminary analysis on a subset of loci and set to 20. Four rounds (two with and two without the outgroups) were performed, to ensure a proper cleaning. Following Tan *et al.* (2015) we only performed a light filtering of alignment positions that contained gaps to reduce signal loss. Sites with more than 75% gaps were removed from the locus alignments using the program seqtools implemented in the package PASTA (Mirarab et al. 2014).

### Exploration of potential bias

To explore potential sources of bias, a correlation analysis between locus properties was performed with the R package Performance Analytics (Peterson and Carl 2018). Then, we explored a possible impact of heterogeneity of evolutionary rates between taxa with three different methods: i) the LS^3^ approach (Rivera-Rivera and Montoya-Burgos 2016, Rivera-Rivera and Montoya-Burgos 2019) ii) a custom approach based on iterative principal component analysis (PCA) of long branch (LB) heterogeneity scores of taxa (Struck 2014) in individual gene trees and iii) different rooting approaches: midpoint rooting, minimal ancestor deviation (MAD) approach (Tria et al. 2017) and minimum variance rooting (MinVar) (Mai et al. 2017). We evaluated a possible impact of base composition heterogeneity among taxa and markers using i) incremental removal of the most GC-biased loci and ii) the custom iterative PCA of GC content of taxa in individual gene trees. For the LS^3^ approach we defined four clades of interest which corresponded to the four highly supported clades recovered in the phylogenetic trees inferred from the mergeR1R2 data set: Clade1= sect. *Pharmacosycea*; Clade2=subg. *Urostigma*, Clade3=sect. *Oreosycea*, Clade4= “gynodioecious clade”. The LS^3^ algorithm was then used to find a subsample of sequences in each locus that evolve at a homogeneous rate across all clades of interests. The minTaxa parameter was set to 1.

We developed a custom iterative PCA approach in R (R Core Team 2018) to analyse LB heterogeneity scores and GC content of the RAD loci. The PCA consisted in the eigenanalysis of the matrix of the correlations between loci and yielded a set of principal axes corresponding to linear combinations of these variables (Manly and Alberto 2017). We used the scores of the taxa along the axes to detect a possible non-random distribution of taxa on the reduced space of the PCA. Depending on the studied properties different groups were highlighted and compared (LB scores: sect. *Pharmacosycea versus* all other fig trees; GC content: *Mixtiflores versus* all other fig trees and then sect. *Pharmacosycea versus* other fig trees, *Mixtiflores* excluded). An initial PCA was performed on all loci and the difference between the groups was statistically assessed by means of a Wilcoxon test applied to the score of the loci upon the first PCA axis. Then, the locus showing the highest correlation with the axis was removed and another PCA and Wilcoxon test were performed on the thinned dataset. The locus showing the highest correlation with the first axis of this new PCA was removed and so on. Only loci for which no structure could be observed were retained for phylogenetic analysis (*i.e* loci for which Wilcoxon tests were non-significant, indicating no difference between the groups along the first axis of the PCA). This approach was implemented to homogenize taxa properties as much as possible prior to phylogenetic inference. PCA were performed with the R package ade4 (Dray and Dufour 2007). The R script and a tutorial to perform iterative PCAs is available from https://github.com/acruaud/radseq_ficus_2020.

### Phylogenetic inference

Gene trees were inferred with a maximum likelihood (ML) approach as implemented in raxmlHPC-PTHREADS-AVX version 8.2.4. A rapid bootstrap search (100 replicates) followed by a thorough ML search (-m GTRGAMMA) was performed. Phylogenetic analyses of the concatenated data set were performed with supermatrix (ML) and coalescent-based summary methods. For the ML approach, we used raxmlHPC-PTHREADS-AVX version 8.2.4 and IQTREE v1.6.7 (Nguyen et al. 2015). Data sets were analysed without partitioning. A rapid bootstrap search (100 replicates) followed by a thorough ML search (-m GTRGAMMA) was implemented for the RAxML approach. IQTREE analysis employed an ML search with the best-fit substitution model automatically selected and branch supports were assessed with ultrafast bootstrap (Minh et al. 2013) and SH-aLRT test (Guindon et al. 2010) (1000 replicates). Trees were annotated with TreeGraph 2.13 (Stöver and Müller 2010) or the R package ape (Paradis et al. 2004).

### Evolution of life history traits

Three key traits (life form, breeding system and pollination mode) were studied. Stochastic mapping (Huelsenbeck et al. 2003) as described in Bollback (2006) and implemented in the R package phytools was utilized to estimate the ancestral state and the number of transitions for each trait. The transition matrix was first sampled from its posterior probability distribution conditioned on the substitution model (10,000 generations of MCMC, sampling every 100 generations). Then, 100 stochastic character histories were simulated conditioned on each sampled value of the transition matrix. Three Markov models were tested: equal rates model (ER) with a single parameter for all transition rates, symmetric model (SYM) in which forward and reverse transition have the same rate and all rates different model (ARD). AIC scores and Akaike weight for each model were computed.

### Computational resources

Analyses were performed on a Dell PowerEdge T630 server with two 10-core Intel(R) Xeon(R) CPUs E5-2687W v3 @ 3.10GHz and on the Genotoul Cluster (INRA, France, Toulouse, http://bioinfo.genotoul.fr/).

## RESULTS

### Molecular phylogenetic inference with outgroup rooting

As a reminder, classifications of the genus *Ficus* are provided in Table 1. Data sets are described in Table 2 and features of taxa are reported in Tables S1-S3. An average of 2*416,834 paired reads; 2,520 ustacks loci and 2,308 cstacks loci were obtained for each sample included in the RAD library (Table S1). In an attempt to solve the tree backbone, we kept only the 583 most conserved loci assembled from forward reads (75% complete matrix, mergeR1 data set), 91% of which (530) were retained in the final (mergeR1R2) data set. Mining of loci in outgroup genomes largely reduced the level of missing data (from *ca* 70-75% in the mergeR1 data set to *ca* 15-30% in the mergeR1R2 data set depending on the outgroup, Table S1). Between 1 and 55 loci were flagged for each sample by Treeshrink (average 15, Table S4). The final data set used for phylogenetic inference (mergeR1R2) was composed of 70 species of *Ficus* and five outgroups. We did not obtain enough reads for *Sparattosyce dioica* to include it in our analysis. Alignment length was 419,945 bp (Table 2). RAxML and IQTREE produced identical topologies with high statistical support (Figure 1, S1; Table 3). Neither gaps (K=0.418, Pvalue=0.342) nor missing data (K=0.384, Pvalue=0.555) were phylogenetically clustered in the ingroup (i.e. taxa with high percentages of missing data /gaps did not cluster together more often than expected by chance). All subgenera except *Ficus* and *Pharmacosycea* were recovered monophyletic with strong support and all non-monospecific sections of the data set except *Ficus* were monophyletic with strong support. Section *Pharmacosycea* was sister to all other *Ficus* species with strong support. The remaining species clustered into two highly supported groups: 1) subg. *Urostigma* and sect. *Oreosycea*; 2) subg. *Ficus, Sycomorus, Sycidium* and *Synoecia*, hereafter named the “gynodioecious clade” for brevity as it clusters all gynodioecious species of fig trees (although a few monoecious species are present in subg. *Sycomorus*). Relationships within the “gynodioecious clade” were strongly supported with subg. *Sycidium* + *F. carica* sister to subg. *Sycomorus* + other species of the subg. *Ficus.* Subsection *Frutescentiae* was sister to a clade grouping sect. *Eriosycea* and subg. *Synoecia*, yet with poor support. We must note that ASTRAL recovered sect. *Oreosycea* sister to the “gynodioecious clade” when the whole mergeR1R2 data set was considered (Figure S1C), though with low support (PP=0.2). However, when sequences with less that 50% locus coverage were removed from each RAD locus, ASTRAL inferred sect. *Oreosycea* sister to subg. *Urostigma* (Figure S2, PP=0.8). Therefore, incomplete locus coverage might have misled individual gene tree inference resulting in this observed switch of position for sect. *Oreosycea* in the coalescence tree. Only two unsupported changes were observed between the ASTRAL and ML trees in the shallowest nodes (within sect. *Conosycea* and *Malvanthera*, Figures S1-S2).

**Table 2.**
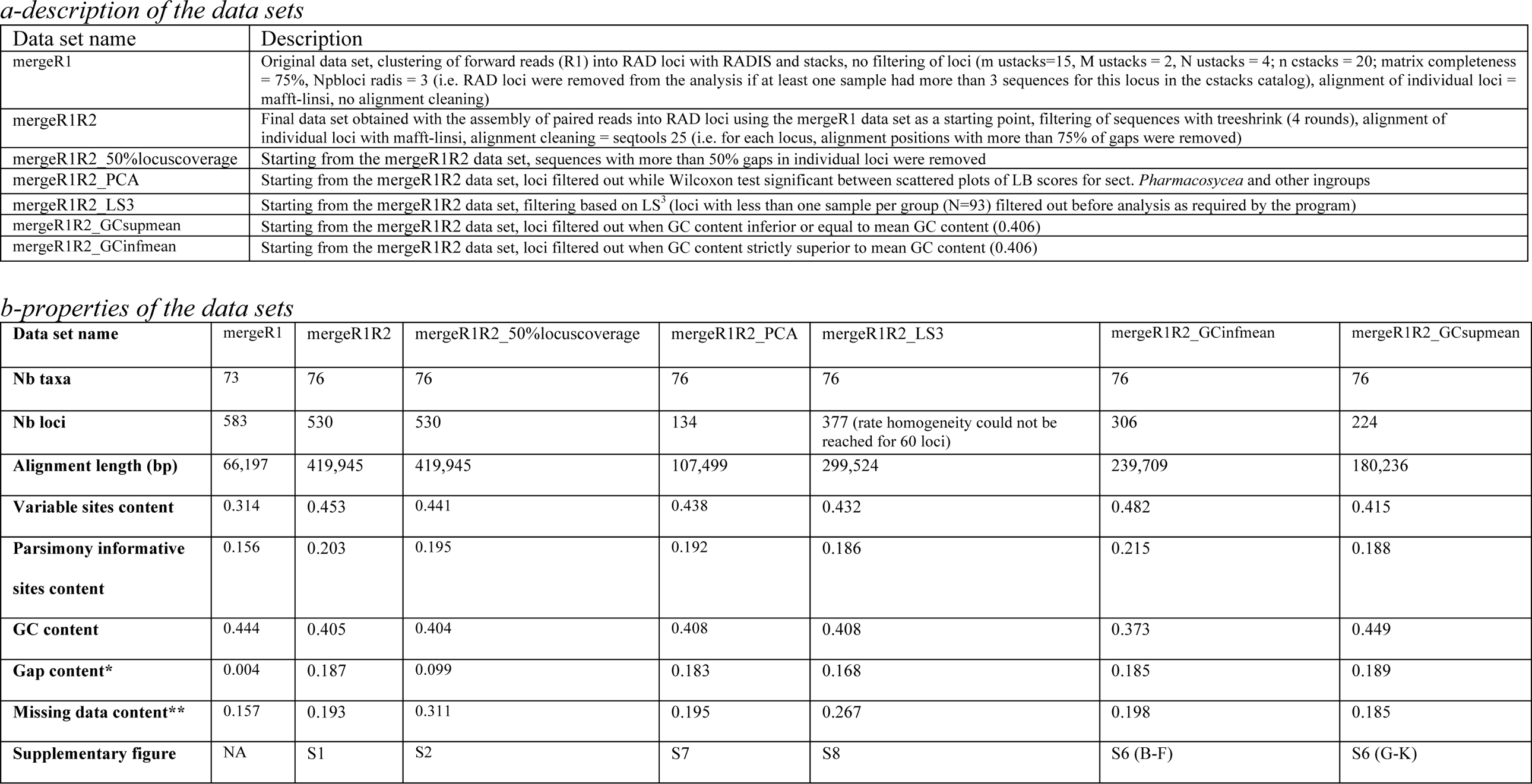
Description of the concatenated data sets analysed in the study. *gap content refers to indels inserted during alignment or missing parts of RAD loci following assembly of forward and reverse reads **missing data refers to missing position in the matrix due to the absence of RAD loci

**Table 3.**
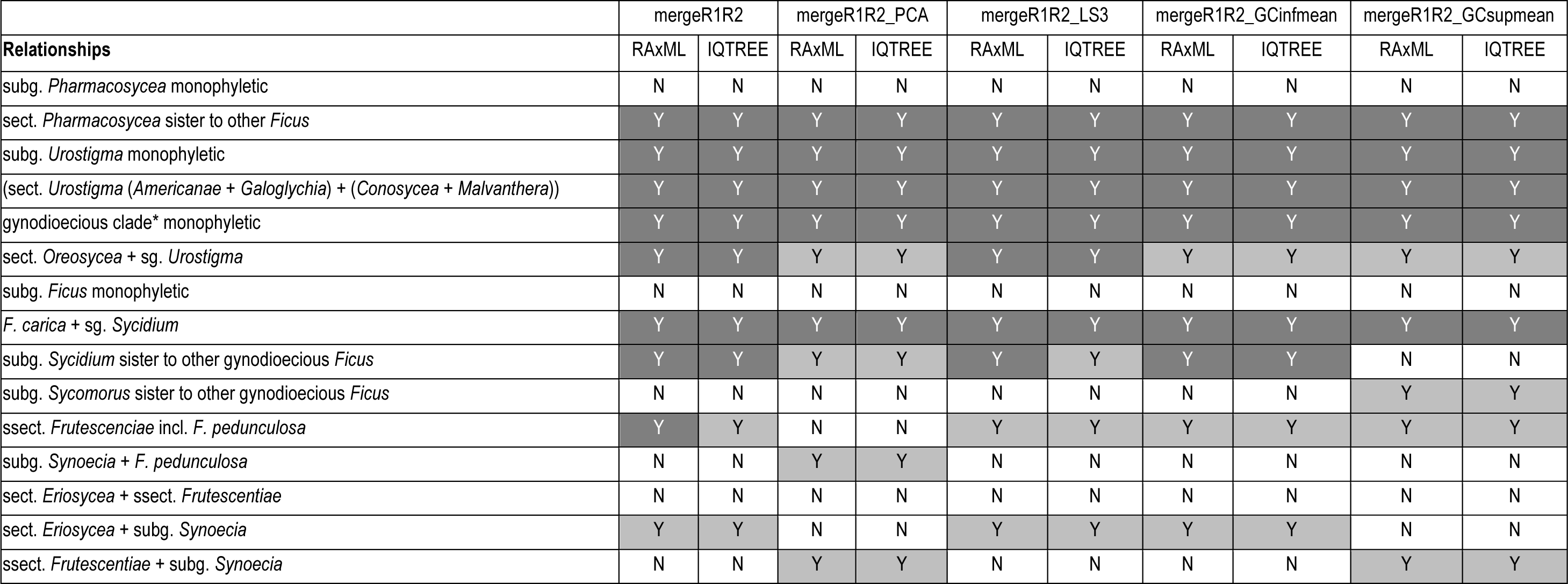
Summary of the phylogenetic relationships inferred from the different data sets when topologies were rooted with outgroups. Data sets are described in Table 2 and topologies are available as supplementary data (Figures S1, S6-8). *Although a few *Sycomorus* are monoecious, we refer to the clade that groups subgenera *Ficus, Sycidium, Sycomorus* and *Synoecia* as the “gynodioecious clade” for brevity. N=Not recovered; Y=recovered. Color coding is as follows: white = not recovered; light grey = RAxML Bootstrap < 90 and IQTREE SH-aLRT < 80 / UFBoot < 95; dark grey = RAxML Bootstrap ≧ 90 and IQTREE SH-aLRT ≧ 80 / UFBoot ≧ 95.

**Figure 1.**
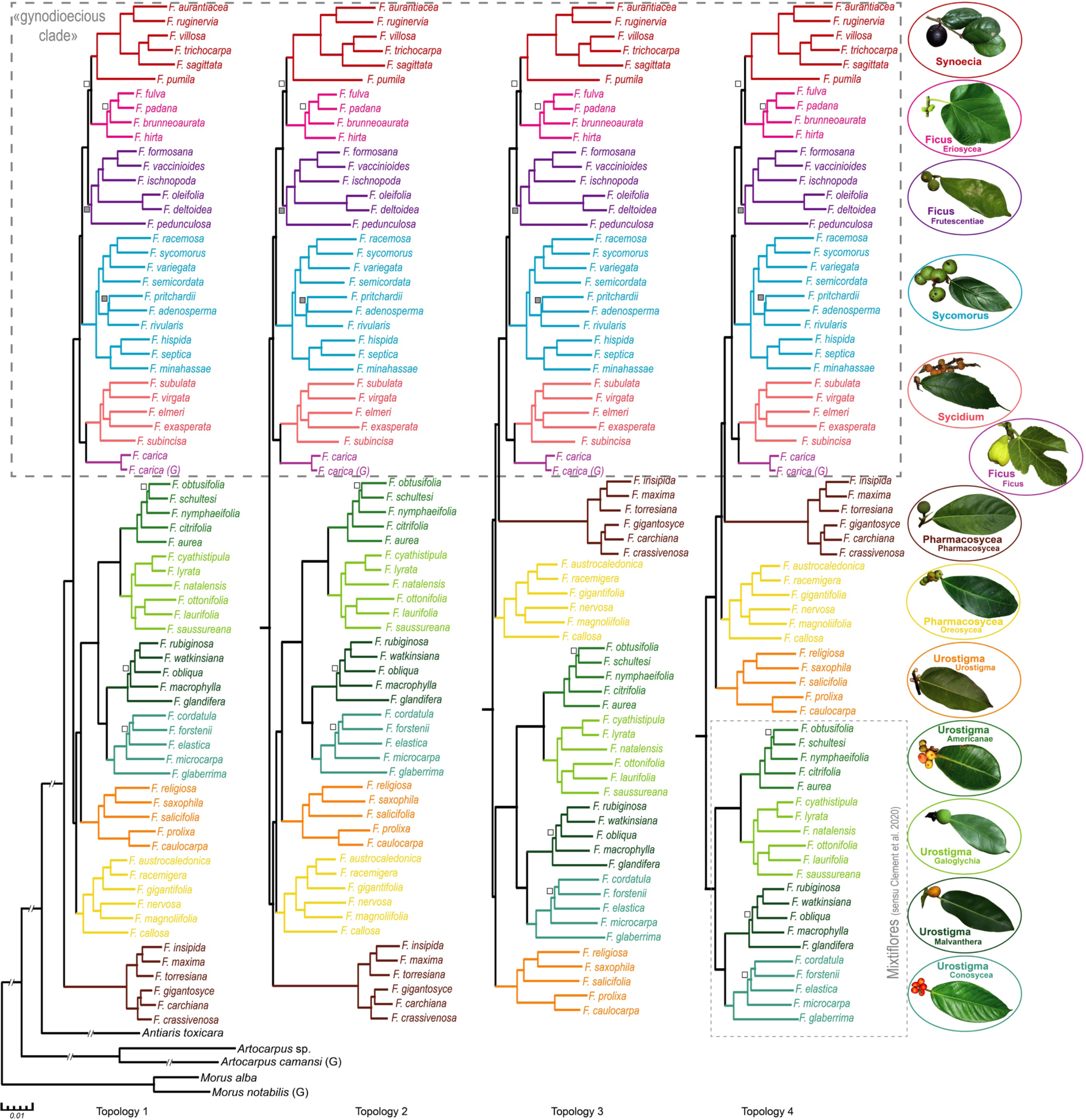
Phylogenetic trees obtained with the molecular data set. RAxML and IQTREE trees were identical. Genomes are indicated with a (G). Subgenera (larger font size) and (sub) sections (smaller font size) are illustrated with pictograms and highlighted with different colors. Classification follows Cruaud *et al.* (2012) as reported in Table 1. All nodes were supported with RAxML Bootstrap (BP) ≧ 90 and IQTREE SH-aLRT ≧ 80 / UFBoot ≧ 95 unless specified with a grey square: BP < 90 or SH-aLRT < 80 / UFBoot < 95 or a white square: BP < 90 and SH-aLRT < 80 / UFBoot < 95. Although a few *Sycomorus* are monoecious, we refer to the clade that groups subgenera *Ficus, Sycidium, Sycomorus* and *Synoecia* as the “gynodioecious clade” for brevity. *Topology 1*: tree obtained from the mergeR1R2 data set with outgroup rooting. This topology is not supported by morphological data (Figures 3 & S9) and supposed to result from an LBA artefact of sect. *Pharmacosycea* to the outgroups. *Topology 2* is obtained for a few data sets when alternative rooting methods are used (Figure 2). It is the most supported by morphological data and evolutionary history of pollinators. *Topology 3* is obtained for a few data sets when alternative rooting methods are used. It is less supported by morphological data and evolutionary history of pollinators than topology 2. *Topology 4* is obtained for a few data sets when alternative rooting methods are used. It is considered unlikely as the position of sect. *Urostigma* is not supported by morphological data. The position of the root could be driven by the GC-content bias exhibited by *Mixtiflores.*

### Identification of potential bias

Spearman’s rank correlation tests showed a significant negative correlation between the proportion of parsimony informative sites or the average bootstrap support of gene trees and i) GC content of loci and ii) LB score heterogeneity of loci (Figure S3). Furthermore, loci with more homogeneous rates among sites (high alpha) were more informative. With the exception of a single unsupported change within subsect. *Eriosycea* (Figure S6), exclusion of GC-rich loci (Table 3) did not induce topological change, which suggested that inferences were not biased by GC content of loci. However, we highlighted compositional bias among samples. As differences in coverage of RAD loci across samples prevented a proper calculation of GC content in the mergeR1R2 data set (i.e. loci were only partially sequenced in some samples), we focused on the mergeR1 data set to explore GC-content bias among samples with PCA (Figure S4). The first principal component (PC1) discriminated between i) all sections of the subg. *Urostigma* except sect. *Urostigma* [*i.e.* the *Mixtiflores* group *sensu* Clement et al. (2020)] and ii) all other species of *Ficus* (eigenvalues = 9.50% for PC1 and 8.57% for PC2, Figure S4Ab). Within the remaining species of *Ficus*, PC1 discriminated between i) sect. *Pharmacosycea* and ii) other species of *Ficus* (eigen-value = 12.55% for PC1 and 6.64% for PC2, Figure S4Ac). Wilcoxon tests showed that the distance separating i) *Mixtiflores* and other fig trees on one side and ii) sect. *Pharmacosycea* and other fig trees (*Mixtiflores* excluded) on the other side was significant for all iterations of the PCA (Figures S4B-C). This means that it was not possible to homogenize GC content of taxa prior to phylogenetic inference to fit with model assumptions. GC content of *Mixtiflores* was significantly higher and GC content of sect. *Pharmacosycea* was significantly lower that GC content of other fig trees (Figures S4D-E).

In addition to heterogeneity of GC content among *Ficus* lineages, we highlighted heterogeneity in evolutionary rates. Two groups were highlighted on the PCA of LB scores across all RAD loci: sect. *Pharmacosycea* and all other fig trees (Figure S5, eigenvalues = 18.76% for PC1 and 4.34% for PC2). Moreover, in average, 29.0% of the loci were flagged for sect. *Pharmacosycea* by LS^3^, while only 19.4% were flagged for other fig trees (Table S5). Attempts to reduce heterogeneity in evolutionary rates with LS^3^ and the custom PCA approach failed. Whatever the concatenated data set analysed (supposedly cleaned or not from bias), the branch leading to sect. *Pharmacosycea* was the longest (Figures 2, S1E, S6D, S6I, S7D, S8C) and sect. *Pharmacosycea* always had significantly higher LB heterogeneity scores than all other taxa (about 7.5 points more for the mergeR1R2_PCA and _LS3 data sets and about 10 points more for the mergeR1R2, _GCinfmean, _GCsupmean data sets; Table 4).

**Table 4.** Comparison of average Long Branch heterogeneity scores (LB scores) between sect. *Pharmacosycea* and all other fig trees in concatenated data sets. Data sets are described in Table 2. Taxa properties are in Table S1. Topologies are available as supplementary data (Figures S1, S6-8)

|  | mergeR1R2 | mergeR1R2_PCA | mergeR1R2_LS3 | mergeR1R2_GCinfmean | mergeR1R2_GCsupmean |
| --- | --- | --- | --- | --- | --- |
| **Sect. *Pharmacosycea*** | -8.39 | -10.58 | -11.40 | -9.58 | -17.75 |
| **Other fig trees** | -18.44 | -18.00 | -18.93 | -19.17 | -6.97 |
| **Wilcoxon test (P-value)** | 6.9e-05 | 0.00018 | 0.00012 | 6.3e-05 | 7.5e-05 |

**Figure 2.**
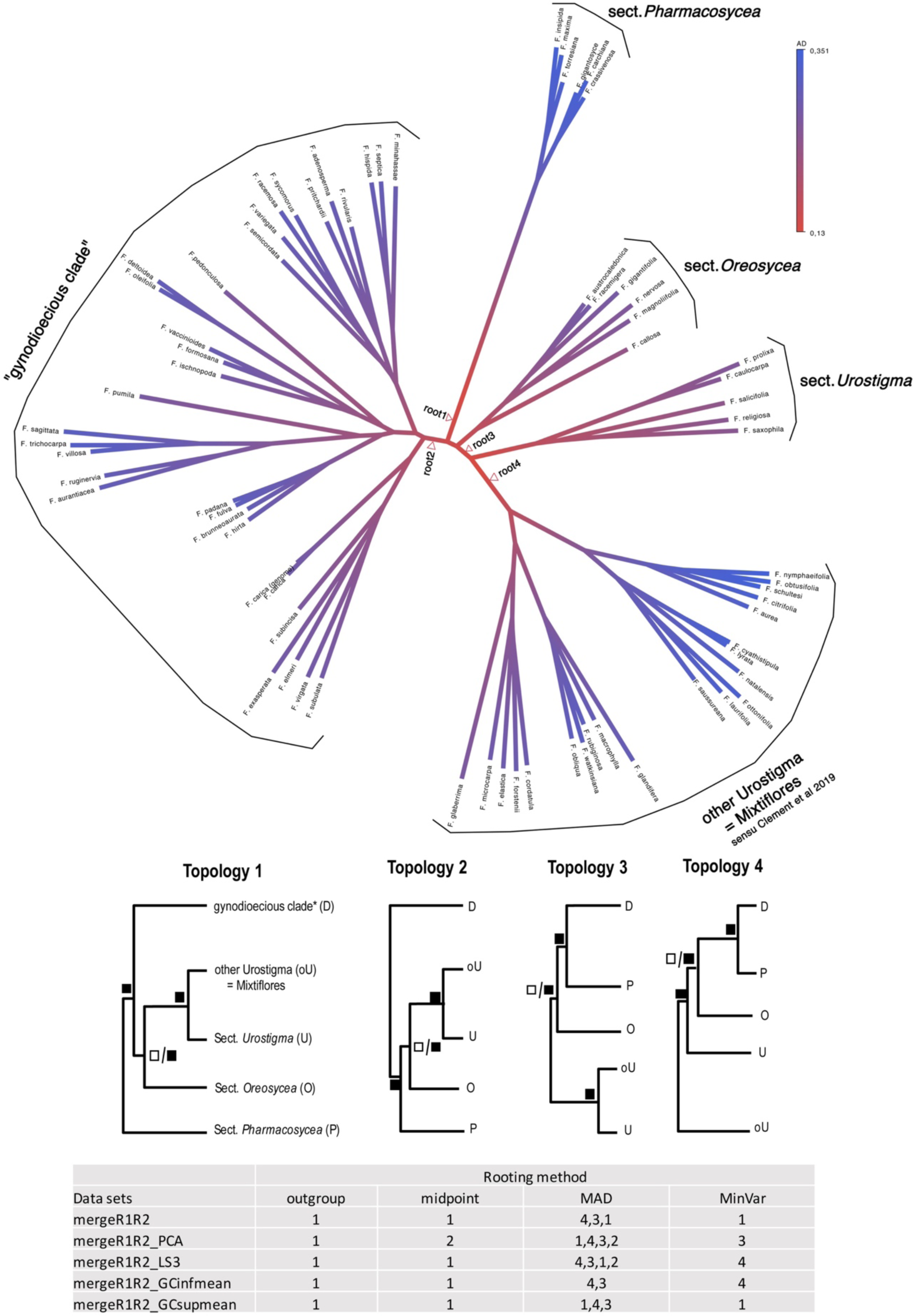
Impact of rooting methods on tree estimation. Three alternative methods to outgroup rooting were tested: midpoint rooting, minimal ancestor deviation (MAD) approach (Tria et al. 2017) and minimum variance rooting (MinVar) (Mai et al. 2017). Outgroups were removed from the data sets described in Table 2 before analysis with IQTREE (best-fit substitution model automatically selected). Full trees are available as supplementary data. *Top figure*: Unrooted tree obtained from the mergeR1R2 data set with branch colors corresponding to their ancestor relative deviation value AD; alternative positions for the root are indicated with arrows. *Middle figure*: Illustration of the four topologies obtained with different rooting strategies. *Bottom figure*: Summary table of the topologies obtained from different data sets and rooting strategies. In all cases, the MAD approach resulted in very high ambiguity scores for the root (0.788-0.999; average=0.938). When alternative positions for the root were identified by MAD the preferred topology is listed first in the table, then, topologies within 0.01 units of ancestral deviation scores are listed by decreasing order of ancestral deviation. Statistical support of nodes varied with analysed data sets. Black squares indicate strong support: IQTREE SH-aLRT ≧ 80 / UFBoot ≧ 95; white squares indicate low support: IQTREE SH-aLRT < 80 / UFBoot < 95. *Although a few *Sycomorus* are monoecious, we refer to the clade that groups subgenera *Ficus, Sycidium, Sycomorus* and *Synoecia* as the “gynodioecious clade” for brevity.

### Impact of rooting strategies on the molecular tree

Fast-evolving or compositionally biased ingroup taxa can be drawn towards the outgroups, especially when the outgroup is distantly related to the ingroup [Long Branch Attraction (LBA) artefact (Bergsten 2005)], which is the case here (Figure S1). For that reason, we tested alternative rooting methods to outgroup rooting. While outgroup rooting always recovered the long-branched sect. *Pharmacosycea* sister to the remaining fig trees (topology 1, Figure 2), other rooting methods suggested three alternative positions for the root: i) on the branch separating the “gynodioecious clade” from other *Ficus* species (topology 2), ii) on the branch separating subg. *Urostigma* from other fig trees (topology 3), iii) on the branch separating sect. *Conosycea, Malvanthera, Americanae* and *Galoglychia* (*Mixtiflores*) from the remaining fig trees (topology 4). The root ambiguity index calculated by MAD was high (0.788-0.999; average=0.938) which indicates that root inference was problematic for all data sets.

### Morphological study

The morphological matrix is provided in Appendix S2. The nexus file that contains the majority-rule consensus tree obtained from the morphological data and the four conflicting RAD topologies on which reconstruction of ancestral character states was performed with Mesquite v3.31 (Maddison and Maddison 2018) can be opened with PAUP* or Mesquite. Among the 102 morphological characters used, 100 were parsimony-informative. The heuristic search yielded 213 equally parsimonious trees of 736 steps long (CI = 0.443, RI = 0.710). The majority-rule consensus (MRC) tree and the strict consensus trees are depicted in Figure 3. The consistency index of each character is provided in Appendix S1. As expected, statistical support was generally low. Only a few nodes, all of them also supported in the molecular tree, received bootstrap supports > 80: sections *Eriosycea* (BP=100), *Malvanthera* (BP=96); *Pharmacosycea* (BP=87) and *Urostigma* (BP=92). Three main clades were recovered: 1) a monophyletic subg. *Pharmacosycea*; 2) subg. *Urostigma*; 3) the “gynodioecious clade”. Interestingly, the MRC tree differed slightly from the outgroup rooted molecular hypothesis (topology 1, Figure 1). The main differences were: 1) the branching order of the most basal nodes with sect. *Oreosycea* + sect. *Pharmacosycea* (BP=54) sister to all other *Ficus* while only sect. *Pharmacosycea* was recovered sister to the remaining fig trees in the molecular topology; 2) the position of *F. carica* that is sister to section *Eriosycea* in the morphological tree *versus* sister to subg. *Sycidium* in the RAD trees; sect. *Urostigma* sister to section *Conosycea* versus sister to all other sections of *Urostigma*; species of subg. *Synoecia* forming a grade within subg. *Ficus*, while it is monophyletic and nested within subg. *Ficus* in the molecular trees. Character transformations inferred with PAUP* on the four competing molecular topologies are illustrated in Figure S9 (ACCTRAN optimization). Topologies 1 and 4 were the less compatible with morphological data; while topology 2 was supported by the highest number of unambiguous transformations followed by topology 3.

**Figure 3.**
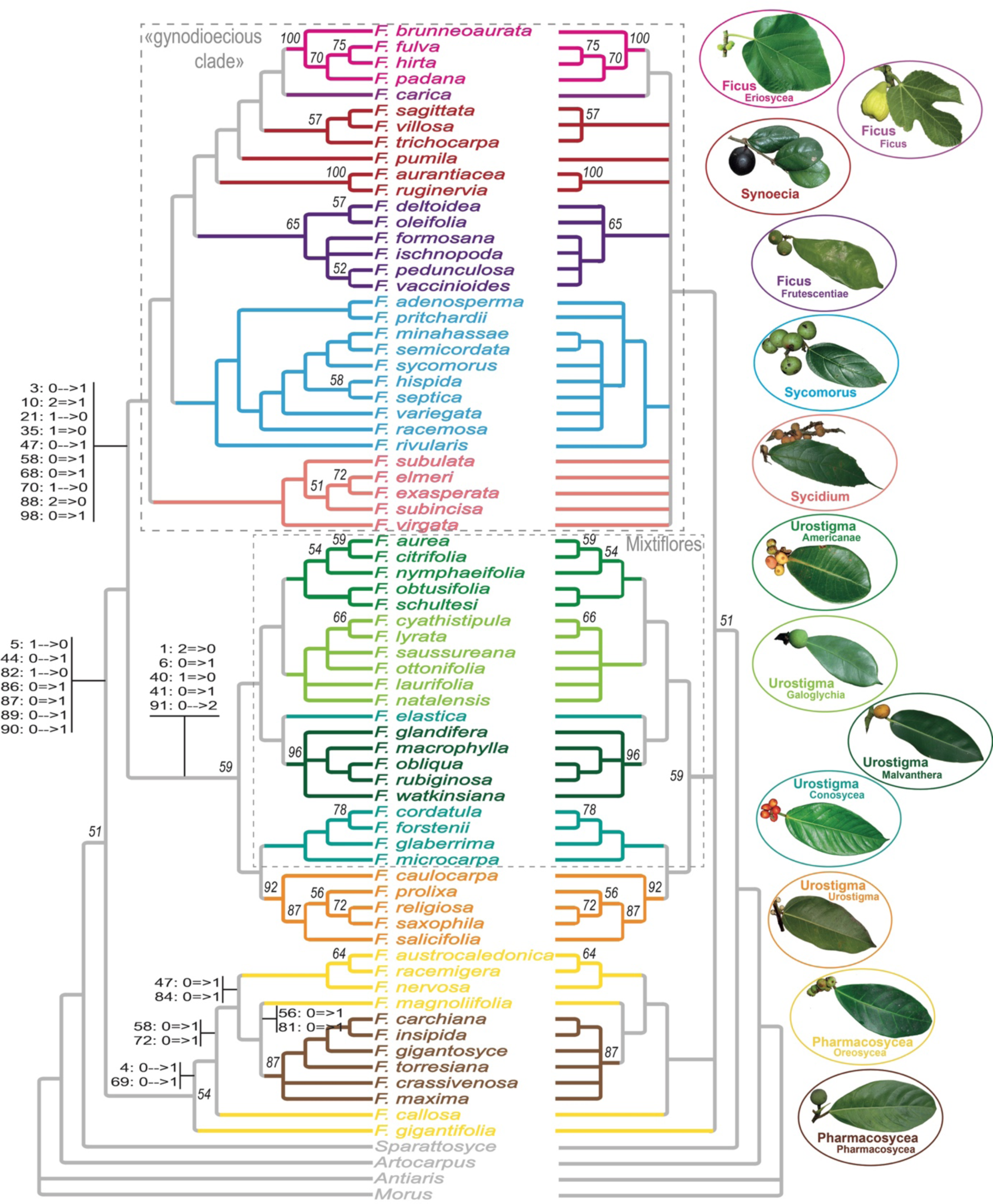
Phylogenetic trees obtained with the morphological data set. *Left*: majority rule consensus tree; *Right*: strict consensus tree. Bootstrap (100 replicates) at nodes. Ambiguous (single arrow) and non-ambiguous (double arrow) transformations inferred by PAUP* (ACCTRAN optimization) are listed for key nodes as follows: character: ancestral state -->/=> derived state (see list of character /states in Appendix S1). Subgenera (larger font size) and (sub)sections (smaller font size) are illustrated with pictograms and highlighted with different colors as for Figure 1. Classification follows Cruaud *et al.* (2012) as reported in Table 1. Although a few *Sycomorus* are monoecious, we refer to the clade that groups subgenera *Ficus, Sycidium, Sycomorus* and *Synoecia* as the “gynodioecious clade” for brevity.

### Reconstruction of traits evolution under stochastic mapping

Because of the low resolution of the morphological tree, reconstructions were performed on the molecular trees only. For all topologies and traits, the ER model had the lowest AIC and highest Akaike weight (Table S6). Therefore, the ER model was subsequently chosen to trace the evolution of breeding system, pollination mode and life form in *Ficus* (Figure S10). Results are given in Table 5, and revealed that the ancestor of all extant *Ficus* was most likely an actively-pollinated, monoecious tree from which hemi-epiphytes /hemi-epilithes and root climbers evolved. Gynodioecy appeared once in the genus and monoecy re-appeared at least twice in the “gynodioecious clade”. Active pollination was lost several times independently.

**Table 5.**
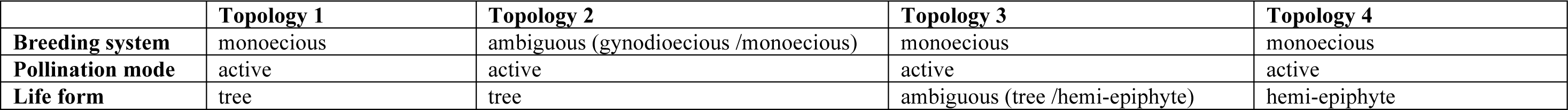
Summary of the reconstruction of traits evolution on the molecular trees. The character state of the most recent common ancestor to all fig trees is mentioned for the three analyzed traits. The full reconstructions can be found in Figure S10. *Nota:* the word hemi-epiphytes is used for all species with aerial roots (i.e. hemi-epiphytes *s.s.* and the few hemi-epilithes).

|  | **Topology 1** | **Topology 2** | **Topology 3** | **Topology 4** |
| --- | --- | --- | --- | --- |
| **Breeding system** | monoecious | ambiguous (gynodioecious /monoecious) | monoecious | monoecious |
| **Pollination mode** | active | active | active | active |
| **Life form** | tree | tree | ambiguous (tree /hemi-epiphyte) | hemi-epiphyte |

## DISCUSSION

### A first phylogenomic hypothesis for fig trees and its morphological counterpart

Here, we propose the first phylogenetic hypothesis for the genus *Ficus* based on pangenomic nuclear markers and a sampling representative of all subgenera and sections (Figure 1). Our analyses highlight heterogeneity in both evolutionary rates and GC content among *Ficus* lineages. We show that sect. *Pharmacosycea* has significantly higher LB heterogeneity scores than all other taxa and, regardless of all the attempts made to reduce this bias (custom PCA approach and LS^3^), the branch leading to sect. *Pharmacosycea* is still, by far, the longest (i.e. branch length /evolutionary rates cannot be properly homogenized) (Table 4, Figures S1, S6-8). In addition, the custom PCA approach shows that it was not possible to homogenize GC content of taxa prior to phylogenetic inference to fit with model assumptions. GC content of *Mixtiflores* (sect. *Americanae, Galoglychia, Conosycea, Malvanthera*) is significantly higher and GC content of sect. *Pharmacosycea* is significantly lower. As heterogeneity in evolutionary rates and base composition are considered important sources of systematic bias (Brinkmann et al. 2005, Philippe et al. 2017), it is crucial to critically interpret results in the light of other line of evidences.

This is why we also built a phylogeny of the same taxa from morphological features. We went one step further than Weiblen (2000) who published the first morphological phylogeny of *Ficus* by increasing the number of characters and taxa analysed. The consistency index values of characters are low (average = 0.375, 32 characters with CI > 0.5), showing a high level of homoplasy as already underlined by Clement & Weiblen (2009) for Moraceae. Nevertheless, the recovered trees are structured enough (Figure 3) to allow comparison with the molecular trees and discuss agreement and discrepancies that could reveal inference bias.

### When next generation sequencing corroborates past generation botanists: agreement between morphological and molecular hypotheses

In agreement with previous molecular studies based on nuclear data, we confirm the monophyly of the subgenera *Sycidium* and *Sycomorus.* The monophyly of *Sycomorus* was not supported in the morphological study by Weiblen (2002) but is confirmed by the morphological analysis presented here. This suggests that the polyphyly of these two subgenera observed by Bruun-Lund *et al.* (2017) in their plastid phylogeny could be due to divergent copies of chloroplastic DNA.

While its monophyly has never been questioned by former botanists (Corner 1958, Berg 1989), the subgenus *Urostigma* had never been recovered as monophyletic in molecular studies so far and one section (*Galoglychia*) was missing from the morphological work by Weiblen (2000) to formally test its monophyly. Here, we highlight a strongly supported subgenus *Urostigma* both with molecular and morphological data. This highly diversified and widespread monoecious subgenus presents a relatively uniform morphology over its range and is well-characterized by non-ambiguous apomorphies (Figure 3): all species have aerial roots and they have only one waxy gland located at the base of the midrib. This subgenus includes, amongst others, sacred banyan trees and giant stranglers.

For the first time in a molecular framework, we highlight a strongly supported clade that groups all gynodioecious fig trees (Figure 1), which corresponds to a previous circumscription of subgenera within *Ficus* [Table 1, (Corner 1958)]. This monophyly was already highlighted in the morphological work by Weiblen (2000) and is confirmed by our morphological analysis (Figure 3). Aside from the breeding system, this clade is well defined by several non-ambiguous synapomorphies on the morphological tree (species have generally less than 10 lateral veins; figs are frequently stipitate; and bears more than three ostiolar bracts; the stigma in short-styled pistillate flowers is mostly cylindrical; and the fruits are compressed).

Finally, and as observed in previous molecular work, the subgenus *Ficus* appeared polyphyletic in our molecular and morphological trees, though results differ between the two approaches as discussed in the next section. It is the first time that the non-monophyly of this subgenus is assessed through phylogenetic inference of morphological data as only two representatives of a single section of the subgenus *Ficus* (sect. *Eriosycea*) were included in previous analysis.

### Systematic bias versus morphological convergences: discrepancies between molecular and morphological evidence

As mentioned above, subgenus *Ficus* appeared polyphyletic in our molecular and morphological trees. On both trees, subg. *Synoecia* is nested within a clade that comprises subsect. *Frutescentiae* and sect. *Eriosycea* of the subg. *Ficus.* On the morphological tree, *F. carica*, the type species of the subg. *Ficus*, is recovered sister to sect. *Eriosycea.* In the molecular tree, *F. carica* does not cluster with other species of the subg. *Ficus* and is instead recovered sister to subg. *Sycidium*. This sister-taxa relationship has already been observed in the past with molecular data, though with low support (Cruaud *et al*., 2012). This result, which is the most surprising result of our study is nevertheless supported by four homoplastic synapomorphies: deciduousness; asymmetrical lamina; margin of perianth hairy and the presence of pistillodes in male flowers, although several species of subg. *Pharmacosycea* and *Sycomorus* also have pistillodes. More species are needed to confirm this result, especially species of the subseries *Albipilae* Corner. However, if the molecular position of *F. carica* is confirmed, then subsect. *Frutescentiae* and sect. *Eriosycea* should not be considered anymore as belonging to subg. *Ficus*.

The evolutionary history of subg. *Synoecia* seems to be linked to the evolutionary history of subg. *Ficus* in its current circumscription. Indeed, in all molecular studies that were representative enough of the biodiversity of the genus, *Synoecia* always clustered with sect. *Eriosycea* and subsect. *Frutescentiae.* In the morphological tree of Weiblen (2000), subg. *Synoecia* appeared monophyletic and sister to sect. *Eriosycea* (no representative of the subsect. *Frutescentiae* was included). The same sister taxa relationship is observed in our molecular tree, though this is the only part of the tree that received low support. *Synoecia* is not recovered as monophyletic in our morphological tree. However, this may be due to the high level of homoplasy in the analysed characters. Again, further studies are needed but *Synoecia* may simply constitute a lineage that has evolved as root climbers as originally suggested by Corner (1965).

The second discrepancy between our morphological and molecular results are the relationships between sect. *Malvanthera, Conosycea* and *Urostigma* of subg. *Urostigma. Malvanthera* and *Conosycea* are recovered sister in all molecular analyses based on nuclear data including ours. However, all *Conosycea* except *F. elastica* are sister to *Urostigma* in our morphological tree. The morphological results may be due to a lack of signal as all species have aerial roots and show a relatively uniform morphology – at least for a set of characters that are meaningful across the entire genus. These observations lead us to consider that the molecular hypotheses better reflect the history of the subgenus *Urostigma*.

The last discrepancy between morphological and molecular data concerns the subgenus *Pharmacosycea*. Sections *Pharmacosycea* and *Oreosycea* form one of the few supported clades of our morphological tree (BP > 50, Figure 3), while they form a grade in the molecular tree (Figure 1). The monophyly of the subgenus *Pharmacosycea* has never been questioned by former botanists (Corner 1958, Berg 1989) but has been challenged by all molecular analyses published so far. It is noteworthy that the morphological strict consensus tree presented by Weiblen (2000) was modified to show a monophyletic sect. *Pharmacosycea* that was present in the majority rule consensus of the bootstrap trees but not in the most parsimonious tree in which *F. albipila* (sect. *Oreosycea*) was sister to *F. insipida* (sect. *Pharmacosycea*) [see legend of Figure 5 and text in Weiblen (2000)]. As for other groups of fig trees, apomorphies shared by sections *Pharmacosycea* and *Oreosycea* are difficult to find because there are always a few species that differ from the original ground plan. However, if we consider unambiguous apomorphies that are shared between sect. *Pharmacosycea* and at least one species of sect. *Oreosycea*, six homoplastic characters can be retained (Figure 3): epidermis of petiole flaking off; absence of colored spot on figs; fig stipe present; staminate flowers scattered among pistillate flowers; pistillode present; pistillate perianth partially connate with tepals fused basally.

It could be argued that morphological results are due to convergence and this argument cannot be definitely ruled out. However, the morphological tree shows a high level of congruence with the molecular tree for other *Ficus* groups and it is difficult understand why morphology would be only misleading for this subgenus. The ecology of the two sections is close but so is the ecology of all species from subg. *Urostigma* for instance. Denser sampling of subg. *Pharmacosycea* in future molecular works, including species of sect. *Oreosycea* subseries *Albipilae* Corner, may help to better resolve relationships between these two groups. While sect. *Pharmacosycea* may not render sect. *Oreosycea* paraphyletic as observed in the morphological tree, they could be at least closely related. On the opposite, exploration of GC content and evolutionary rates clearly show that sect. *Pharmacosycea* does not exhibit the same properties as all other fig trees, which could mislead molecular inferences.

### Where fig trees take root: rooting issues, higher-level relationships and clues from the pollinator tree of life

Indeed, deeper phylogenetic relationships remain the most problematic issue. The unrooted molecular tree highlights the problem we face to resolve the root of fig trees (Figure 2): 1) a long branch leading to a recent diversification of sect. *Pharmacosycea*, and 2) short surrounding branches supporting the “gynodioecious clade”, sect. *Oreosycea* and subg. *Urostigma*, suggesting fast diversification of the ancestors of the present lineages. This pattern is predicted to favor artefactual rooting of trees when distant outgroups are used. Further, the recently developed minimal ancestor deviation (MAD) approach that is robust to variation in evolutionary rates among lineages (Tria et al. 2017) shows that root inference is problematic in the original data set (mergeR1R2) and in all other data sets built to test for potential bias (Figure 2).

Four competing topologies are suggested. Given the long branch leading to sect. *Pharmacosycea* and the impossibility to homogenize its evolutionary rate (and GC content) with those of other fig trees, it seems reasonable to suspect that Topology 1 results from an LBA artefact (Bergsten 2005). Indeed, long-branched taxa can cluster with outgroups with high statistical support irrespective of their true phylogenetic relationships (convergent changes along the two long branches are interpreted as false synapomorphies because current models do not reflect evolutionary reality) (Phillips et al. 2004). It is known that LBA tend to be reinforced as more and more data are considered (e.g. (Boussau et al. 2014), which is probably the case here, even though we used an outgroup belonging to Castillae, *i.e.* the closest relative of *Ficus* (Clement and Weiblen 2009). So far, in all molecular studies, sect. *Pharmacosycea* was recovered sister to all other fig trees [with low or high support, but see Figure 1 of Clement *et al.* (2019), which depicts the phylogeny of Involucraoideae, where section *Albipilae* was sister to all other fig trees with low support. A result that is not recovered from the data set centered on *Ficus* spp. on their Figure 2]. Topology 1 contradicts morphological evidences (Figure 3) and previous classification (Table 1). We must note that the unbalance between overall short internal branches and long branches leading to sect. *Pharmacosycea* on one side and outgroups on the other is recurrently observed in all molecular-based analyses of the *Ficus* phylogeny. Hence, all molecular analyses might have been trapped by an LBA artefact with a misplacement of the root on the long branch leading to sect. *Pharmacosycea*. There is a last line of evidence that should be considered to critically interpret the phylogenies presented here: the evolutionary history of the pollinators. Interestingly, the position of the pollinators of sect. *Pharmacosycea* (genus *Tetrapus*) as sister to all other species of Agaonidae recovered in early molecular studies has been shown to result from an LBA artefact (Cruaud et al. 2012). In addition, pollinators of sect. *Oreosycea* (*Dolichoris*) and sect. *Pharmacosycea* (*Tetrapus*) are also grouped by morphology (see supplementary data of Cruaud *et al.* 2012). Notably *Dolichoris* and *Tetrapus* share a unique metanotal structure in males of agaonids (Figures S14A & B in Cruaud et al., 2012). This structural character advocates close relationships instead of convergence between their host figs.

Although they cannot be used as direct evidence, these results are converging evidence suggesting that topology 1 does not accurately reflect early divergence events within fig trees. There are three known techniques to reduce LBA (Bergsten 2005). The first possibility is reducing the branch length of the ingroup taxa that are drawn towards the outgroups. Here, we show that the branch leading to sect. *Pharmacosycea* was still significantly longest regardless of the attempt made to reduce this bias. The second possibility is outgroup removal (Bergsten 2005). However, as sect. *Pharmacosycea* is among the first lineages to diverge this method is not helpful. The third possibility is increasing sampling. This should definitely be attempted in the future but without guarantee as LBA may be too strong to be broken (Boussau et al. 2014). More generally, given strong bias highlighted here, increasing species sampling appears at least as relevant (if not more) as increasing the number of sequenced regions for each species analysed.

Topology 4 is the least supported by morphological data (Figure S9) and appears unlikely given the highly supported monophyly of subg. *Urostigma* in morphological analyses (Figure 3). A possible explanation for its recovery would be the high GC content of *Mixtiflores* that could be linked to a high number of SNPs supporting the group and an increased branch length, which distorts the calculation of ancestral deviation. In addition, given life forms observed in Moraceae if appears most likely that the ancestor of fig trees was a freestanding tree, which contradicts the scenario of trait evolution based on topology 4 (Table 5, Figure S10).

Therefore, two topologies still remain likely (topologies 2 & 3, Figure 1). They only differ by the position of the grade composed by sections *Oreosycea* and *Pharmacosycea*. In topology 2, these sections cluster with subg. *Urostigma* and the genus *Ficus* is divided into two groups: monoecious and gynodioecious species. In topology 3, the sections cluster with the “gynodioecious clade” and the genus is split into species with aerial roots on one side and other fig trees on the other side. Topology 2 is the most supported by morphological data on fig trees (Figure S9). Importantly, unambiguous transformations that support the monoecious /gynodioecious split are not only linked to breeding system (or pollination mode). Indeed, characters supporting the split are related to tree height; number of lateral veins; fig stipitate or not; pistillate flowers sessile or not; shape of stigma in short-styled pistillate flowers; lamina thickness; lamina margin; furcation of lateral veins; tertiary venation; presence of interfloral bracts. On the opposite, topology 3 and, more specifically, the sister taxa relationships between sect. *Oreosycea*, sect. *Pharmacosycea* and the “gynodioecious clade” is not supported by any unambiguous morphological transformation (Figure S9).

Finally, the current molecular phylogeny of pollinators of fig trees provide more support to topology 2. Indeed, and again even though this argument should be used only as an indirect evidence, reconciliation of the agaonid tree of life with topology 3 would require a partial reversal of the pollinator tree that contradicts transformation series for the structure of male mesosoma and female basal flagellomeres (anelli) (see Cruaud *et al.* 2012, Figure S14). The current pollinator tree supports a progressive fusion of male mesosoma and female anelli, while topology 3 would favor a subdivision of mesosoma and basal flagellomere as the most derived character state, a trend that has never been observed in Hymenoptera.

Therefore, after considering all the evidence (bias, morphology, pollinators) we consider that the most likely topology for the *Ficus* tree of life is topology 2. Interestingly, this topology agrees with one of the first statement of Corner (1958): “the first division of *Ficus* is into the monoecious and dioecious species, but it is more convenient to recognize three subgenera, namely the monoecious banyans (*Urostigma*), the monoecious trees (*Pharmacosycea*) and the dioecious *Ficus*”.

Nevertheless, increasing sampling is required to resolve the root of the *Ficus* tree of life. Until now, no investigation was undertaken to identify another root for *Ficus* than the one identified by outgroup rooting. However, resolving this issue is important not only to understand the evolution of *Ficus*, but is also crucial to our understanding of key questions such as how species’ traits have changed over time and how fig trees co-diversified with their pollinating wasps. Here we go one step further in our understanding of the evolution of key traits as previous trees were not enough resolved to allow inference. While there is a consensus concerning the ancestral pollination mode that is inferred as active for all topologies, ambiguity remains for ancestral breeding system (including for our favored topology, Figure 4). Finally, it appears most likely that the ancestor of fig trees was a freestanding tree.

**Figure 4.**
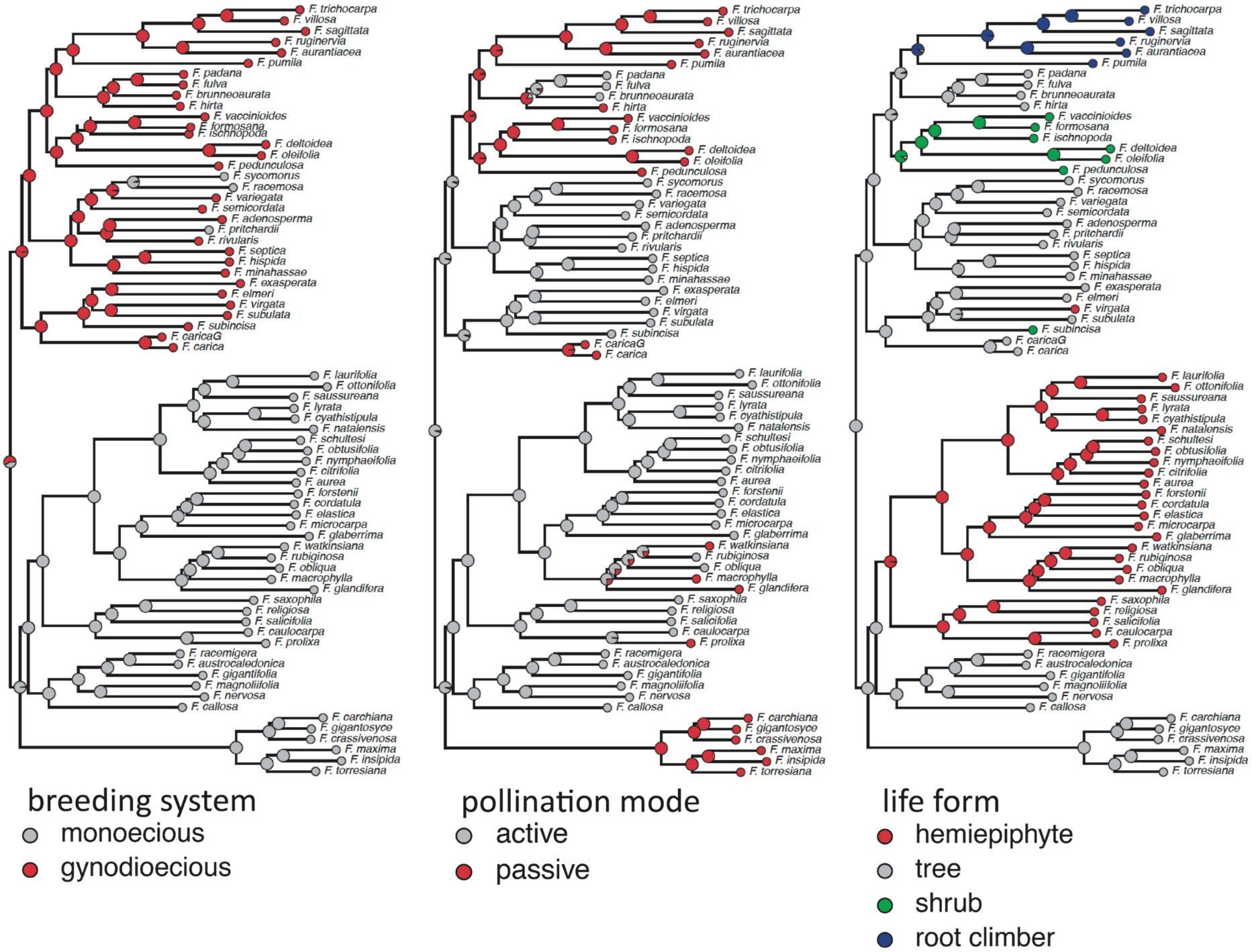
Reconstruction of traits evolution on the favored topology (topology 2). All reconstructions are available in Figure S10. *Nota:* the word hemi-epiphytes is used for all species with aerial roots (i.e. hemi-epiphytes *s.s.* and the few hemi-epilithes).

## CONCLUSION

We present the first phylogenetic hypothesis for the genus *Ficus* based on pangenomic nuclear markers and a sampling effort representative of all subgenera and sections. For the first time, we recover a monophyletic and strongly supported subgenus *Urostigma* and a strongly supported clade grouping all gynodioecious fig trees. Our in-depth analysis of biases, general pattern of rooting preferences and morphological data completed with indirect evidence from the pollinator tree of life, highlights that previous molecular studies might have been trapped by LBA, which resulted in an artefactual placement of sect. *Pharmacosycea* as sister to all other fig trees. The next key step will be to increase sample size. Indeed, confidence in phylogenetic inference may increase with increasing data sets encompassing as much as possible the overall diversity in the studied group. Increasing species sampling appears at least as relevant as increasing the number of sequenced regions for each species analysed. This work is a first step towards a clarification of the classification and evolutionary history of fig trees. Taxonomic changes foreseen by previous molecular works and new ones will need to be undertaken. But taxonomists will need to be cautious and humble as invalidating current and widely used names will generate more confusion than clarity.

## ACKNOWLEDGEMENTS

We thank S. Santoni (INRAE) for technical advices, MGX for sequencing, the Genotoul bioinformatics platform Toulouse Midi-Pyrenees for computing resources and C.C. Berg and E.J.H. Corner for their eternal contributions. This work was partially funded by UPD-OVCRD (171715 PhDIA) and UPD-NSRI (BIO-18-1-02). AC thanks C. J. Rivera-Rivera for his help with running LS^3^.

## AUTHOR CONTRIBUTIONS

J.Y.R. and A.C. conceived the study, analyzed data and drafted the manuscript. J.Y.R, L.J.R, Y.Q.P, D.R.Y, F.K., A.B, R.D.H., R.U., R.A.S.P, S.F, A.C. collected samples. J.Y.R identified or verified identification of all samples. L.J.R., L.S. C.T.C. and A.C. performed lab work. M.G and J.P.R contributed scripts. All authors gave final approval for publication.

## DATA ACCESSIBILITY

Raw paired reads are available as an NCBI Sequence Read Archive (#SRPXXXX). Pipeline and scripts are available from: https://github.com/acruaud/radseq_ficus_2020. Data sets and trees are available from the Dryad Digital Repository: http://dx.doi.org/10.5061/dryad.[NNNN]

